# Active inference under visuo-proprioceptive conflict: Simulation and empirical results

**DOI:** 10.1101/795419

**Authors:** Jakub Limanowski, Karl Friston

**Affiliations:** Wellcome Centre for Human Neuroimaging, Institute of Neurology, University College London, London, UK

**Keywords:** action, active inference, attention, body representation, multisensory integration, precision, predictive coding

## Abstract

It has been suggested that the brain controls hand movements via internal models that rely on visual and proprioceptive cues about the state of the hand. In active inference formulations of such models, the relative influence of each modality on action and perception is determined by how precise (reliable) it is expected to be. The ‘top-down’ affordance of expected precision to a particular sensory modality is associated with attention. Here, we asked whether increasing attention to (i.e., the precision of) vision or proprioception would enhance performance in a hand-target phase matching task, in which visual and proprioceptive cues about hand posture were incongruent. We show that in a simple simulated agent—based on predictive coding formulations of active inference—increasing the expected precision of vision or proprioception improved task performance (target matching with the seen or felt hand, respectively) under visuo-proprioceptive conflict. Moreover, we show that this formulation captured the behaviour and self-reported attentional allocation of human participants performing the same task in a virtual reality environment. Together, our results show that selective attention can balance the impact of (conflicting) visual and proprioceptive cues on action—rendering attention a key mechanism for a flexible body representation for action.

## Introduction

Controlling the body’s actions in a constantly changing environment is one of the most important tasks of the human brain. The brain solves the complex computational problems inherent in this task by using internal probabilistic (Bayes-optimal) models^1-6^. These models allow the brain to flexibly estimate the state of the body and the consequences of its movement, despite noise and conduction delays in the sensorimotor apparatus, via iterative updating by sensory prediction errors from multiple sources. The state of the hand, in particular, can be informed by vision and proprioception. Here, the brain makes use of an optimal integration of visual and proprioceptive signals, where the relative influence of each modality—on the final ‘multisensory’ estimate—is determined by its relative reliability or precision, depending on the current context^6-15^.

These processes can be investigated under an experimentally induced conflict between visual and proprioceptive information. The underlying rationale here is that incongruent visuo-proprioceptive cues about hand position or posture have to be integrated (provided the incongruence stays within reasonable limits), because the brain’s body model entails a strong prior belief that information from both modalities is generated by one and the same external cause; namely, one’s hand. Thus, a partial recalibration of one’s unseen hand position towards the position of a (fake or mirror-displaced) hand seen in an incongruent position has been interpreted as suggesting an (attempted) resolution of visuo-proprioceptive conflict to maintain a prior body representation^16-23^.

Importantly, spatial or temporal perturbations can be introduced to visual movement feedback *during action*—by displacing the seen hand position in space or time, using video recordings or virtual reality. Such experiments suggest that people are surprisingly good at adapting their movements to these kinds of perturbations; i.e., they adjust to novel visuo-motor mappings by means of visuo-proprioceptive recalibration or adaptation^9,10,24-26^. During motor tasks involving the resolution of a visuo-proprioceptive conflict, one typically observes increased activity in visual and multisensory brain areas^27-30^. The remapping required for this resolution is thought to be augmented by attenuation of proprioceptive cues^10,24,26,31^. The conclusion generally drawn from these results is that visuo-proprioceptive recalibration (or visuo-motor adaptation) relies on temporarily adjusting the weighting of conflicting visual and proprioceptive information to enable adaptive action under specific prior beliefs about one’s ‘body model’.

The above findings and their interpretation can in principle be accommodated within a hierarchical predictive coding formulation of active inference as a form of Bayes-optimal motor control, in which proprioceptive *as well as* visual prediction errors can update higher-level beliefs about the state of the body and thus influence action^32-34^. Hierarchical predictive coding rests on a probabilistic mapping from unobservable causes (hidden states) to observable consequences (sensory states), as described by a hierarchical generative model, where each level of the model encodes conditional expectations (*‘beliefs’*) about states of the world that best explains states of affairs encoded at lower levels (i.e., sensory input). The causes of sensations are inferred via model inversion, where the model’s beliefs are updated to accommodate or ‘explain away’ ascending prediction error (a.k.a. Bayesian filtering or predictive coding^35-37^). Active inference extends hierarchical predictive coding from the sensory to the motor domain, in that the agent is now able to fulfil its model predictions via *action*^38^. In brief, movement occurs because high-level multi- or amodal beliefs about state transitions predict proprioceptive and exteroceptive (visual) states that would ensue if a particular movement (e.g. a grasp) was performed. Prediction error is then suppressed throughout the motor hierarchy^3,39^, ultimately by spinal reflex arcs that enact the predicted movement. This also implicitly minimises exteroceptive prediction error; e.g. the predicted visual action consequences^34,40-42^. Crucially, all ascending prediction errors are precision-weighted based on model predictions (where precision corresponds to the inverse variance), so that a prediction error that is expected to be more precise has a stronger impact on belief updating. The ‘top-down’ affordance of precision has been associated with *attention*^43-45^.

This suggests a fundamental implication of attention for behaviour, as action should be more strongly informed by prediction errors ‘selected’ by attention. In other words, the impact of visual or proprioceptive prediction errors on multisensory beliefs driving action should not only depend on factors like sensory noise, but may also be regulated via the ‘top-down’ affordance of precision; i.e., by directing the focus of selective attention towards one or the other modality.

Here, we used a predictive coding scheme^6,38^ to test this assumption. We simulated behaviour (i.e., prototypical grasping movements) under active inference, in a simple hand-target phase matching task (Fig. 1) during which conflicting visual or proprioceptive cues had to be prioritized. Crucially, we included a condition in which proprioception had to be adjusted to maintain visual task performance and a converse condition, in which proprioceptive task performance had to be maintained in the face of conflicting visual information. This enabled us to address the effects reported in the visuo-motor adaptation studies reviewed above and studies showing automatic biasing of one’s own movement execution by incongruent action observation^46,47^. In our simulations, we asked whether changing the relative precision afforded to vision versus proprioception—corresponding to attention—would improve task performance (i.e., target matching with the respective instructed modality, vision or proprioception) in each case. We implemented this ‘attentional’ manipulation by adjusting the inferred precision of each modality, thus changing the degree with which the respective prediction errors drove model updating and action. We then compared the results of our simulation with the actual behaviour and subjective ratings of attentional focus of healthy participants performing the same task in a virtual reality environment. We anticipated that participants, in order to comply with the task instructions, would adopt an ‘attentional set’^48-50^ prioritizing the respective instructed target tracking modality over the task-irrelevant one^51-53^, by means of internal precision adjustment—as evaluated in our simulations.

**Figure 1.**
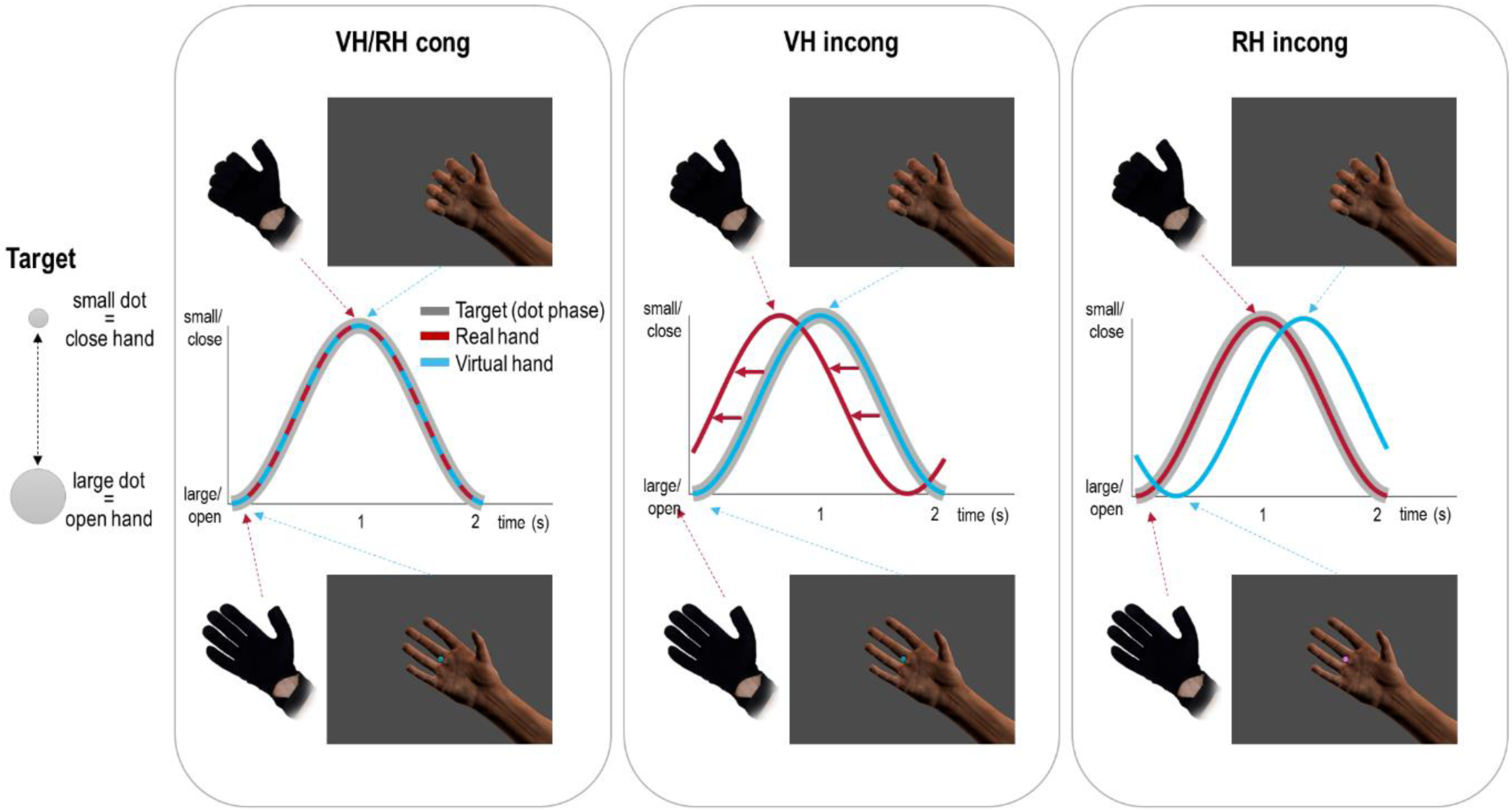
Task design and behavioural requirements. We used the same task design in the simulated and behavioural experiments, focusing on the effects of attentional modulation on hand-target phase matching via prototypical (i.e., well-trained prior to the experiment, see Methods) oscillatory grasping movements at 0.5 Hz. Participants (or the simulated agent) controlled a virtual hand model (seen on a computer screen) via a data glove worn on their unseen right hand. The virtual hand (VH) therefore represented seen hand posture (i.e., vision), which could be uncoupled from the real hand posture (RH; i.e., proprioception) by introducing a temporal delay (see below). The task required matching the phase of one’s right-hand grasping movements to the oscillatory phase of the fixation dot (‘target’), which was shrinking-and-growing sinusoidally at 0.5 Hz. In other words, participants had to rhythmically close the hand when the dot shrunk and to open it when the dot expanded. Our design was a balanced 2 × 2 factorial design: The task was completed (or simulated) under congruent or incongruent hand movements: the latter were implemented by adding a lag of 500 ms to the virtual hand movements (Factor ‘congruence’). Furthermore, the participants (or the simulated agent) performed the task with one of two goals in mind: to match the movements of the virtual hand (VH) or of those of the real hand (RH) to the phase of the dot (Factor ‘instructed modality’; written instructions were presented before each trial, and additionally represented by the fixation dot’s colour). Note that whereas in the congruent conditions (VH *cong*, RH *cong*) both hand positions were identical, and therefore both hands’ grasping movements could simultaneously be matched to the target’s oscillatory phase (i.e., the fixation dot’s size change), only one of the hands’ (virtual or real) movements could be phase-matched to the target in the incongruent conditions—necessarily implying a phase mismatch of the other hand’s movements. In the VH *incong* condition, participants had to adjust their movements to counteract the visual lag; i.e., they had to phase-match the virtual hand’s movements (i.e., vision) to the target by shifting their real hand’s movements (i.e., proprioception) out of phase with the target. Conversely, in the RH *incong* condition, participants had to match their real hand’s movements (i.e., proprioception) to the target’s oscillation, and therefore had to ignore the fact that the virtual hand (i.e., vision) was out of phase. The curves show the performance of an ideal participant (or simulated agent).

## Results

### Simulation results

We based our simulations on predictive coding formulations of active inference^6,13,38,45^. In brief (please see Methods for details), we simulated a simple agent that entertained a generative model of its environment (i.e., the task environment and its hand), while receiving visual and proprioceptive cues about hand posture (and the target). Crucially, the agent could act on the environment (i.e., move its hand), and thus was engaged in active inference.

The simulated agent had to match the phasic size change of a central fixation dot (target) with the grasping movements of the unseen real hand (proprioceptive hand information) or the seen virtual hand (visual hand information). Under visuo-proprioceptive conflict (i.e., a phase shift between virtual and real hand movements introduced via temporal delay), only one of the hands could be matched to the target’s oscillatory phase (see Fig. 1 for a detailed task description). The aim of our simulations was to test whether—in the above manual phase matching task under perceived visuo-proprioceptive conflicts—increasing the expected precision of sensory prediction errors from the instructed modality (vision or proprioception) would improve performance, whereas increasing the precision of prediction errors from the ‘distractor’ modality would subvert performance. Such a result would demonstrate that—in an active inference scheme—behaviour under visuo-proprioceptive conflict can be augmented via top-down precision control; i.e., selective attention^43-45^. In our predictive coding-based simulations, we were able to test this hypothesis by changing the precision afforded to prediction error signals— related to visual and proprioceptive cues about hand posture—in the agent’s generative model.

Figures 2-3 show the results of these simulations, in which the ‘active inference’ agent performed the target matching task under the two kinds of instruction (virtual hand or real hand task; i.e., the agent had a strong prior belief that the visual or proprioceptive hand posture would track the target’s oscillatory size change) under congruent or incongruent visuo-proprioceptive mappings (i.e., where incongruence was realized by temporally delaying the virtual hand’s movements with respect to the real hand). In this setup, the virtual hand corresponds to hidden states generating visual input, while the real hand generates proprioceptive input.

**Figure 2.**
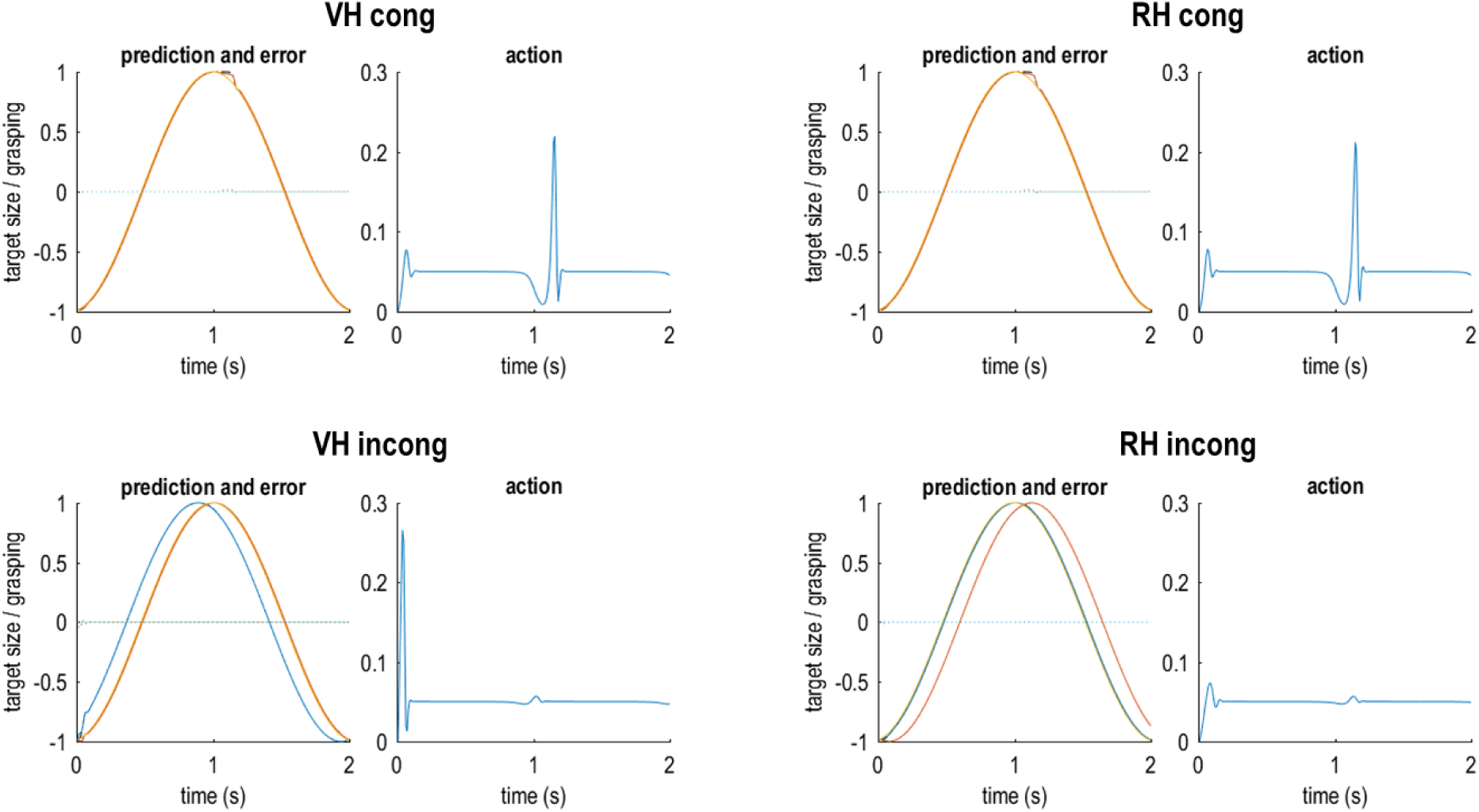
Simulated behaviour of an agent performing the hand-target phase matching task under ideally adjusted model beliefs. Each pair of plots shows the simulation results for an agent with a priori ‘ideally’ adjusted (but unrealistic, see text for details) model beliefs about visuo-proprioceptive congruence; i.e., in the congruent tasks, the agent believed that its real hand generated matching seen and felt postures, whereas it believed that the same hand generated mismatching postures in the incongruent tasks. Note that these simulations are unrealistic in that the agent would not perceive visuo-proprioceptive conflicts at all. Each pair of plots shows the simulation results for one grasping movement in the VH and RH tasks under congruence or incongruence; the left plot shows the predicted sensory input (solid coloured lines; yellow = target, red = vision, blue = proprioception) and the true, real-world values (broken black lines) for the target and the visual and proprioceptive hand posture, alongside the respective sensory prediction errors (dotted coloured lines; blue = target, green = vision, purple = proprioception); the right plot (blue line) shows the agent’s action (i.e., the rate of change in hand posture, see Methods). Note that target phase matching is near perfect and there is practically no sensory prediction error (i.e., the dotted lines stay around 0).

**Figure 3.**
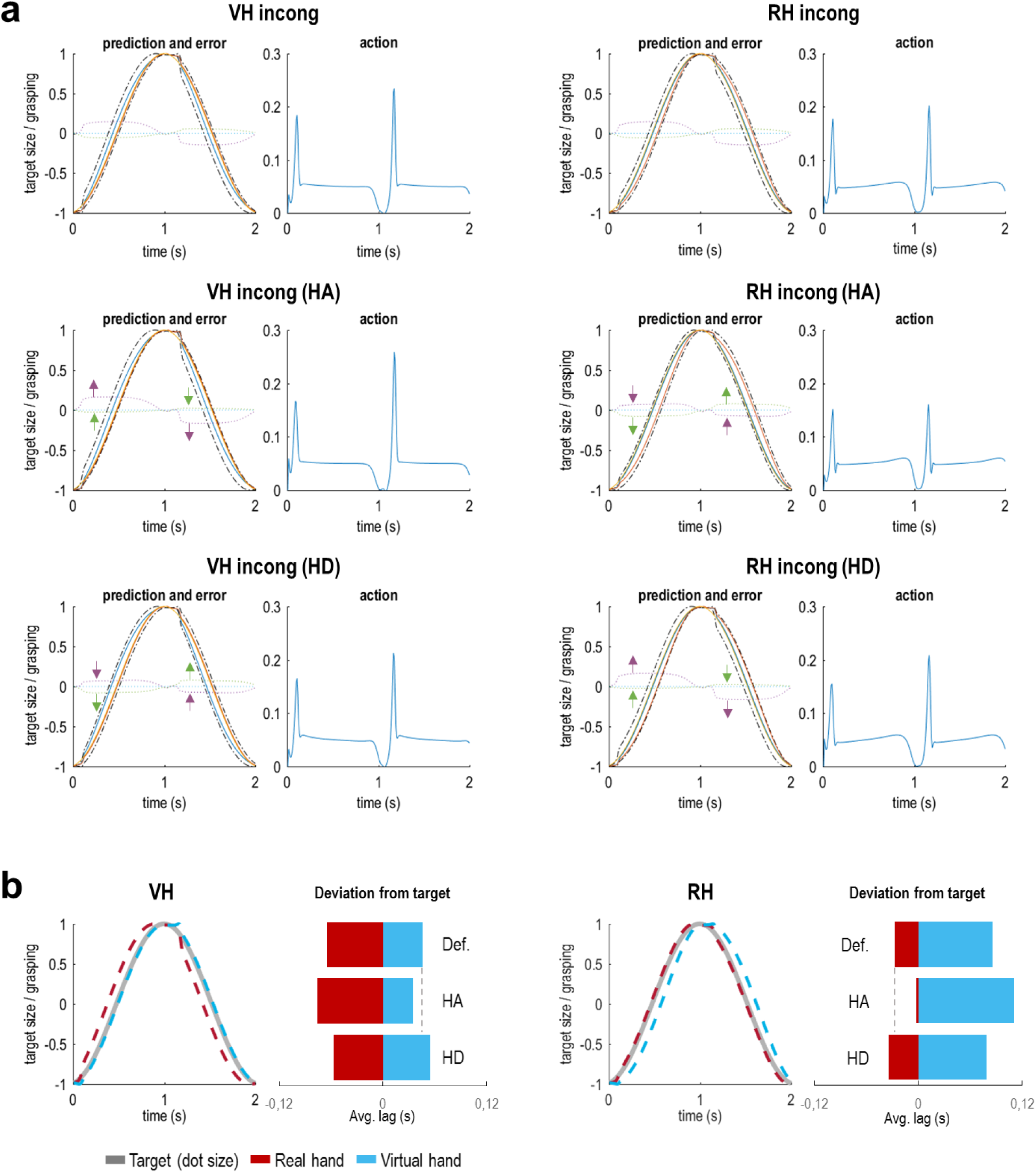
Simulated behaviour of a ‘realistic’ agent performing the hand-target phase matching task. Here we simulated an agent performing the incongruent tasks under the prior belief that its hand generated matching visual and proprioceptive information; i.e., under perceived visuo-proprioceptive conflict. **(a)** The plots follow the same format as in Fig. 2. Note that, in these results, one can see a clear divergence of true from predicted visual and proprioceptive postures, and correspondingly increased prediction errors. The top row shows the simulation results for the default weighting of visual and proprioceptive information; the middle row shows the same agent’s behaviour when precision of the respective task-relevant modality (i.e., vision in the VH task and proprioception in the RH task) was increased (HA: high attention); the bottom row shows the analogous results when the precision of the respective other, irrelevant modality was increased (HD: high distraction). Note how in each case, increasing (or decreasing) the log precision of vision or proprioception resulted in an attenuation (or enhancement) of the associated prediction errors (indicated by green and purple arrows for vision and proprioception, respectively). Crucially, these ‘attentional’ effects had an impact on task performance, as evident by an improved hand-target tracking with vision or proprioception, respectively. This is shown in panel **(b):** The curves show the tracking in the HA conditions. The bar plots represent the average deviation (phase shift or lag, in seconds) of the real hand’s (red) or the virtual hand’s (blue) grasping movements from the target’s oscillatory size change in each of the simulations shown in panel (a). Note that under incongruence (i.e., a constant delay of vision), reducing the phase shift of one modality always implied increasing the phase shift of the other modality (reflected by a shift of red *and* blue bars representing the average proprioceptive and visual phase shift, respectively). Crucially, in both RH and VH incong conditions, increasing attention (HA; i.e., in terms of predictive coding: the precision afforded to the respective prediction errors) to the task-relevant modality enhanced task performance (relative to the default setting, Def.), as evident by a reduced phase shift of the respective modality from the target phase. The converse effect was observed when the agent was ‘distracted’ (HD) by paying attention to the respective task-irrelevant modality.

Under congruent mapping (i.e., in the absence of visuo-proprioceptive conflict) the simulated agent showed near perfect tracking performance (Fig. 2). We next simulated an agent performing the task under incongruent mapping, while equipped with the prior belief that its seen and felt hand postures were in fact unrelated, i.e., never matched. Not surprisingly, the agent easily followed the task instructions and again showed near perfect tracking with vision or proprioception, under incongruence (Fig. 2). However, as noted above, it is reasonable to assume that human participants would have the strong prior belief—based upon life-long learning and association—that their manual actions generated matching seen and felt postures (i.e., a prior belief that modality specific sensory consequences have a common cause). Our study design assumed that this association would be very hard to update, and that consequently performance could only be altered via adjusting expected precision of vision vs proprioception (see Methods).

Therefore, we next simulated the behaviour (during the incongruent tasks) of an agent embodying a prior belief that visual and proprioceptive cues about hand state were in fact *congruent*. As shown in Fig. 3A, this introduced notable inconsistencies between the agent’s model predictions and the true states of vision and proprioception, resulting in elevated prediction error signals. The agent was still able to follow the task instructions, i.e., to keep the (instructed) virtual or real hand more closely matched to the target’s oscillatory phase, but showed a drop in performance compared with the ‘idealized’ agent (cf. Fig. 2).

We then simulated the effect of our experimental manipulation, i.e., of increasing precision of sensory prediction errors from the respective task-relevant (constituting increased attention) or task-irrelevant (constituting increased distraction) modality on task performance. We expected this manipulation to affect behaviour; namely, by how strongly the respective prediction errors would impact model belief updating and subsequent performance (i.e., action).

The results of these simulations (Fig. 3a) showed that increasing the precision of vision or proprioception—the respective instructed tracking modality—resulted in reduced visual or proprioceptive prediction errors. This can be explained by the fact that these ‘attended’ prediction errors were now more strongly accommodated by model belief updating (about hand posture). Conversely, one can see a complementary increase of prediction errors from the ‘unattended’ modality. The key result, however, was that the above ‘attentional’ alterations substantially influenced hand-target phase matching performance (Fig. 3b). Thus, increasing the precision of the instructed task-relevant sensory modality’s prediction errors led to improved target tracking (i.e. a reduced phase shift of the instructed modality’s grasping movements from the target’s phase). In other words, if the agent attended to the instructed visual (or proprioceptive) cues more strongly, its movements were driven more strongly by vision (or proprioception)—which helped it to track the target’s oscillatory phase with the respective modality’s grasping movements. Conversely, increasing the precision of the ‘irrelevant’ (not instructed) modality in each case impaired tracking performance.

The simulations also showed that the amount of action itself was comparable across conditions (blue plots in Figs. 2-3; i.e., movement of the hand around the mean stationary value of 0.05), which means that the kinematics of the hand movement per se were not biased by attention. Action was particularly evident in the initiation phase of the movement and after reversal of movement direction (open-to-close). At the point of reversal of movement direction, conversely, there was a moment of stagnation; i.e., changes in hand state were temporarily suspended (with action nearly returning to zero). In our simulated agent, this briefly increased uncertainty about hand state (i.e., which direction the hand was moving), resulting in a slight lag before the agent picked up its movement again, which one can see reflected by a small ‘bump’ in the true hand states (Figs. 2-3). These effects were somewhat more pronounced during movement under visuo-proprioceptive incongruence and prior belief in congruence—which indicates that the fluency of action depended on sensory uncertainty.

In sum, these results show that the attentional effects of the sort we hoped to see can be recovered using a simple active inference scheme; in that precision control determined the influence of separate sensory modalities—each of which was generated by the same cause, i.e., the same hand—on behaviour by biasing action towards cues from that modality.

### Empirical results

Participants practiced and performed the same task as in the simulations (please see Methods for details). We first analysed the post-experiment questionnaire ratings of our participants (Fig. 4) to the following two questions: “How difficult did you find the task to perform in the following conditions?” (Q1, answered on a 7-point visual analogue scale from “very easy” to “very difficult”) and “On which hand did you focus your attention while performing the task?” (Q2, answered on a 7-point visual analogue scale from “I focused on my real hand” to “I focused on the virtual hand”). For the ratings of Q1, a Friedman’s test revealed a significant difference between conditions (*χ*^2^ _(3,69)_= 47.19, *p* < 0.001). Post-hoc comparisons using Wilcoxon’s signed rank test showed that, as expected, participants reported finding both tasks more difficult under visuo-proprioceptive incongruence (VH *incong* > VH *cong, z*_(23)_ = 4.14, *p* < 0.001; RH *incong* > RH *cong, z*_(23)_ = 3.13, *p* < 0.01). There was no significant difference in reported difficulty between VH *cong* and RH *cong*, but the VH *incong* condition was perceived as significantly more difficult than the RH *incong* condition (*z*_(23)_ = 2.52, *p* < 0.05). These results suggest that, per default, the virtual hand and the real hand instructions were perceived as equally difficult to comply with, and that in both cases the added incongruence increased task difficulty—more strongly so when (artificially shifted) vision needed to be aligned with the target’s phase.

**Figure 4.**
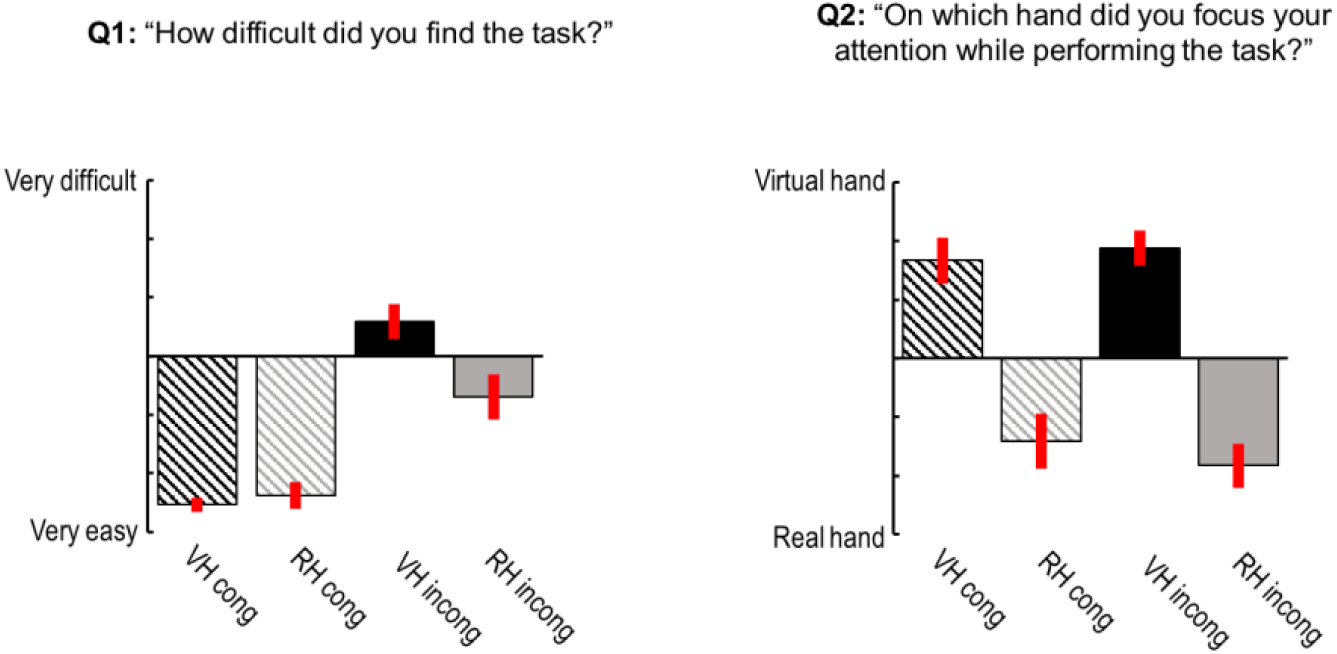
Self-reports of task difficulty and attentional focus given by our participants. The bar plots show the mean ratings for Q1 and Q2 (given on a 7-point visual analogue scale), with associated standard errors of the mean. On average, participants found the VH and RH task more difficult under visuo-proprioceptive incongruence—more strongly so when artificially shifted vision needed to be aligned with the target’s phase (VH incong, Q1). Importantly, the average ratings of Q2 showed that participants attended to the instructed modality (irrespective of whether the movements of the virtual hand and the real hand were congruent or incongruent).

For the ratings of Q2, a Friedman’s test revealed a significant difference between conditions (*χ*^2^(3,69) = 35.83, *p* < 0.001). Post-hoc comparisons using Wilcoxon’s signed rank test showed that, as expected, participants focussed more strongly on the virtual hand during the virtual hand task and more strongly on the real hand during the real hand task. This was the case for congruent (VH *cong* > RH *cong, z*_(23)_ = 3.65, *p* < 0.001) and incongruent (VH *incong* > RH *incong, z*_(23)_ = 4.03, *p* < 0.001) movement trials. There were no significant differences between VH *cong* vs VH *incong*, and RH *cong* vs RH *incong*, respectively. These results show that participants focused their attention on the instructed target modality, irrespective of whether the current movement block was congruent or incongruent. This supports our assumption that participants would adopt a specific attentional set to prioritize the instructed target modality.

Next, we analysed the task performance of our participants; i.e., how well the virtual (or real) hand’s grasping movements were phase-matched to the target’s oscillation (i.e., the fixation dot’s size change) in each condition. Note that under incongruence, better target phase-matching with the virtual hand implies a worse alignment of the real hand’s phase with the target, and *vice versa*. We expected (cf. Fig. 1; confirmed by the simulation results, Figs. 2-3) an interaction between task and congruence: participants should show a better target phase-matching of the virtual hand under visuo-proprioceptive incongruence, if the virtual hand was the instructed target modality (but no such difference should be significant in the congruent movement trials, since virtual and real hand movements were identical in these trials). All of our participants were well trained (see Methods), therefore our task focused on average performance benefits from attention (rather than learning or adaptation effects).

The participants’ average tracking performance is shown in Figure 5. A repeated-measures ANOVA on virtual hand-target phase-matching revealed significant main effects of task (*F*_(1,22)_ = 31.69, *p* < 0.001) and congruence (*F*_(1,22)_ = 173.42, *p* < 0.001) and, more importantly, a significant interaction between task and congruence (*F*_(1,22)_ = 50.69, *p* < 0.001). Post-hoc *t*-tests confirmed that there was no significant difference between the VH *cong* and RH *cong* conditions (*t*_(23)_ = 1.19, *p* = 0.25), but a significant difference between the VH *incong* and RH *incong* conditions (*t*_(23)_ = 6.59, *p* < 0.001). In other words, in incongruent conditions participants aligned the phase of the virtual hand’s movements significantly better with the dot’s phasic size change when given the ‘virtual hand’ than the ‘real hand’ instruction. Furthermore, while the phase shift of the *real* hand’s movements was larger during VH *incong* > VH *cong* (*t*_(23)_ = 9.37, *p* < 0.001)—corresponding to the smaller phase shift, and therefore better target phase-matching, of the virtual hand in these conditions—participants also exhibited a significantly larger shift of their real hand’s movements during RH *incong* > RH *cong* (*t*_(23)_ = 4.31, *p* < 0.001). Together, these results show that participants allocated their attentional resources to the respective instructed modality (vision or proprioception), and that this was accompanied by significantly better target tracking in each case—as expected based on the active inference formulation, and as suggested by the simulation results.

**Figure 5.**
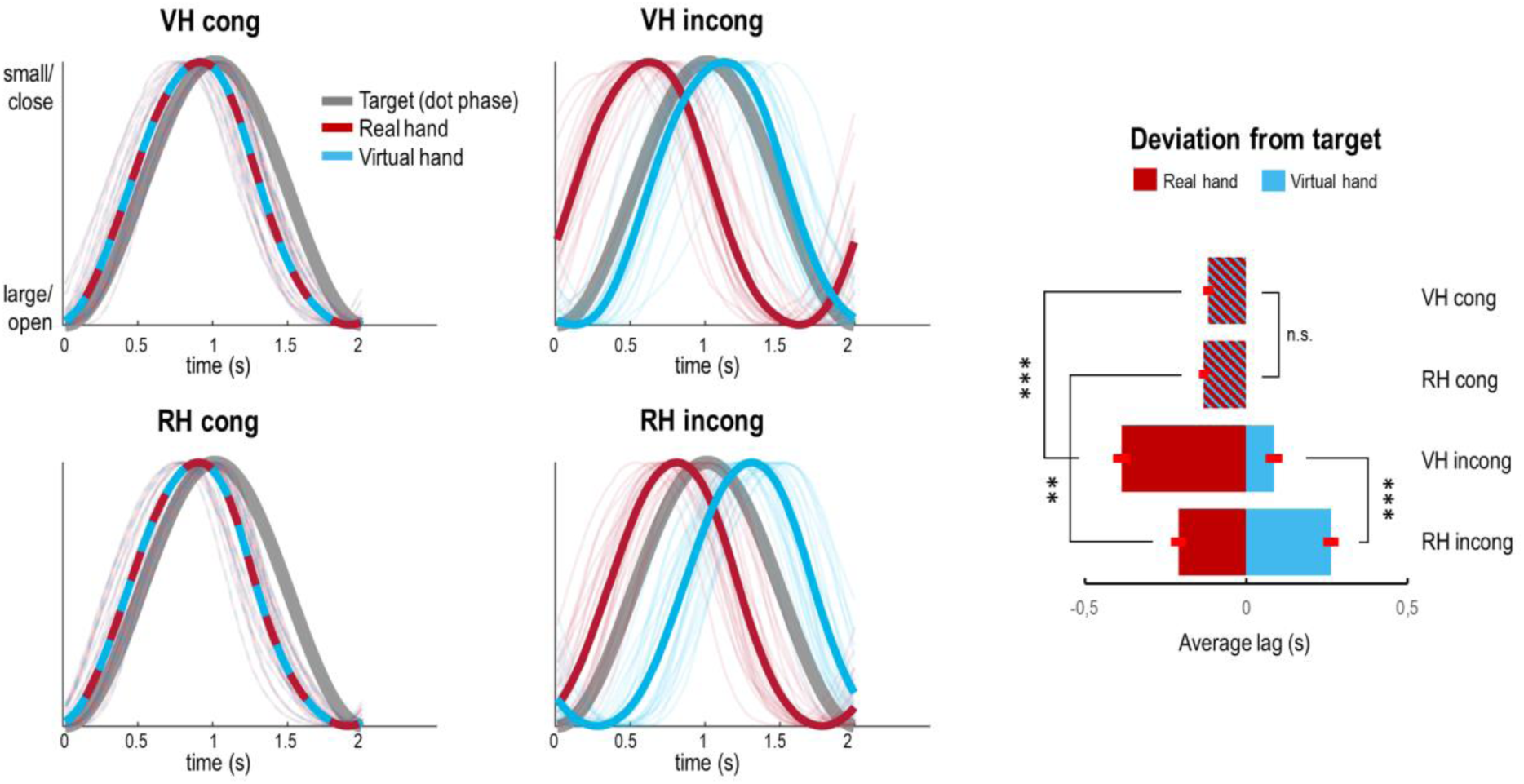
Task performance of our participants. Left: Averaged and normalized trajectories of the real hand’s (red) and the virtual hand’s (blue) grasping movements relative to the oscillation of the target (pulsating fixation dot, grey) in each condition. The individual participant’s averaged and normalized trajectories in each condition are shown as thin lines. In the congruent conditions, the virtual hand’s and the real hand’s movements were identical, whereas the virtual hand’s movements were delayed by 500 ms in the incongruent conditions. Right: The bar plot shows the corresponding average deviation (lag in seconds) of the real hand (red) and the virtual hand (blue) from the target in each condition, with associated standard errors of the mean. Crucially, there was a significant interaction effect between task and congruence; participants aligned the virtual hand’s movements better with the target’s oscillation in the VH *incong* > RH *incong* condition (and correspondingly, the real hand’s movements in the RH *incong* > VH *incong* condition), in the absence of a significant difference between the congruent conditions. Bonferroni-corrected significance: \*\**p* < 0.01, \*\*\**p* < 0.001.

## Discussion

We have shown that behaviour in a hand-target phase matching task, under visuo-proprioceptive conflict, benefits from adjusting the balance of visual versus proprioceptive precision; i.e., increasing attention to the respective task-relevant modality. Our results generally support a predictive coding formulation of active inference, where visual and proprioceptive cues affect multimodal beliefs that drive action—depending on the relative precision afforded to each modality^6,45^. Firstly, a simulated agent exhibited better hand-target phase matching when the expected precision of the instructed ‘task-relevant’ modality (i.e., attention to vision or proprioception) was increased relative to the ‘task-irrelevant’ modality. This effect was reversed when attention was increased to the ‘task-irrelevant’ modality, effectively corresponding to cross-modal distraction. These results suggest that more precise sensory prediction errors have a greater impact on belief updating—which in turn guides goal-directed action. Our simulations also suggest that the effects of changing precision were related to a perceived visuo-proprioceptive conflict—based on a prior belief that one’s hand movements should generate matching visual and proprioceptive sensations. In an agent holding the unrealistic belief that visual and proprioceptive postures were per default unrelated, no evidence for an influence of visuo-proprioceptive conflict on target tracking was observed. Secondly, the self-report ratings of attentional allocation and the behaviour exhibited by human participants performing the same task, in a virtual reality environment, suggest an analogous mechanism: Our participants reported shifting their attention to the respective instructed modality (vision or proprioception)—and they were able to correspondingly align either vision or proprioception with an abstract target (oscillatory phase) under visuo-proprioceptive conflict.

A noteworthy result of the behavioural experiment was a more pronounced shift of real hand movements in the ‘real hand’ condition (i.e., participants partly aligned the delayed virtual hand with the target’s phase). This behaviour resembled that of our simulated agent under ‘high distraction’; i.e., under increased precision of task-irrelevant visual hand cues. This suggests that, in the RH *incong* condition, participants may have been distracted by (attending to) vision. Interestingly, however, our participants reported attentional focus on their real hand and even found the ‘real hand’ task easier than the ‘virtual hand’ task under visuo-proprioceptive incongruence. This suggests that they did not *notice* their ‘incorrect’ behavioural adjustment. One interpretation of this seemingly ‘automatic’ visual bias is suggested by predictive coding formulations of shared body representation and self-other distinction; namely, the balance between visual and proprioceptive prediction errors to disambiguate between ‘I am observing an action’ or ‘I am moving’^3,13^. Generally, visual prediction errors have to be attenuated during action observation to prevent actually realising the observed movement (i.e., mirroring)^13^. However, several studies have demonstrated ‘automatic’ imitative tendencies during action observation, reminiscent of ‘echopraxia’, which are extremely hard to inhibit. For example, seeing an incongruent finger or arm movement biases movement execution^46,47^. In a predictive coding framework, this can be formalized as an ‘automatic’ update of multimodal beliefs driving action by precise (i.e., not sufficiently attenuated) visual body information^3,13,30^. Such an interpretation would be in line with speculations that participants in visuo-motor conflict tasks attend to vision, rather than proprioception, if not instructed otherwise^48,51,53,54^. Our simulation results suggest that altered precision expectations may mediate these effects.

Our results offer new insights into the multisensory mechanisms of a body representation for action, complementing existing theoretical and empirical work. Generally, our results support the notion that an endogenous ‘attentional set’^48-50^ can influence the precision afforded to vision or proprioception during action, and thus prioritize either modality for the current behavioural context. Several studies have shown that visuo-proprioceptive recalibration is context dependent; in that either vision or proprioception may be the ‘dominant’ modality—with corresponding recalibration of the ‘non-dominant’ modality^9,10,48,51-53,55-57^. Thus, our results lend tentative support to arguments that visuo-proprioceptive (or visuo-motor) adaptation and recalibration can be enhanced by increasing the precision of visual information (attending to vision^48,51^). Notably, our results also suggest that the reverse can be true; namely, visuo-proprioceptive recalibration can be counteracted by attending to proprioception. In other words, our results suggest that updating the predictions of a ‘body model’ affects goal-directed action. Crucially, the qualitative similarity of simulated and empirical behaviour provides a mechanistic explanation for these processes, which is compatible with a neurobiological implementation (i.e., predictive coding).

Previous work on causal inference suggests that Bayes-optimal cue integration can explain a variety of multisensory phenomena under intersensory conflict; including the recalibration of the less precise modality onto the more precise one^7,14,15,58-63^. However, in the context of multisensory integration for body (upper limb) representation, the focus of previous models was on perceptual (causal) inference^59,62,64^ or on adaptation or learning^2,65^. Our work advances on these findings by showing that adjusting the precision of two conflicting sources of bodily information (i.e., seen or felt hand posture, which were expected to be congruent based on fixed prior model beliefs in a common cause) enhances the accuracy of *goal-directed action* (i.e., target tracking) with the respective ‘attended’ modality. By allowing our agent to move and optimising sensory precision, this work goes beyond modelling *perceptual* (causal) inference to consider *active* inference; where the consequences of action affect perceptual inference and *vice versa*. Specifically, we showed that *action* (in our case, hand-target tracking) was influenced by *attentional allocation* (to visual or proprioceptive cues about hand position), via augmentation of the impact of sensory prediction errors on model belief updating from the ‘attended’ modality relative to the ‘unattended’ one. In other words, we showed an interaction between sensory attention and (instructed) behavioural goals in a design that allowed the agent (or participant) to actively change sensory stimuli. In short, we were able to model the optimisation of precision—that underwrites multisensory integration—and relate this to sensory attention and attenuation during action. These results generalise previous formulations of sensorimotor control^64,65^ to address attentional effects on action.

The relevance of our model—and the simulation results—also stems from the fact that it is based on a ‘first principles’ approach that, in contrast to most work in this area, commits to a neurobiologically plausible implementation scheme; i.e., predictive coding^35-37^. The model can therefore be thought of as describing recurrent message passing between hierarchical (cortical) levels to suppress prediction error. Model beliefs, prediction errors, and their precision can therefore be associated with the activity of specific cell populations (deep and superficial pyramidal cells; see Methods)^6,40,45^. This means that, unlike most normative models, the current model can, in principle, be validated in relation to evoked neuronal responses—as has been demonstrated in simulations of oculomotor control using the same implementation of active inference^66,67^.

There are some limitations of the present study that should be addressed by future work. Firstly, our task design focuses on prototypical movements and average phase matching. Our results should therefore be validated by designs using more complicated movements. The main aim of our simulations was to provide a proof-of-concept that attentional effects during visuo-motor conflict tasks were emergent properties of active inference formulation. Therefore, our simulations are simplified approximations to a complex movement paradigm with a limited range of movements and deviations. This simplified setup allowed us to sidestep a detailed consideration of forward or generative models for the motor plant—to focus on precision or gain control. In the future, we plan to fit generative models to raw complicated movements. The aim here is to develop increasingly realistic generative models, whose inversion will be consistent with known anatomy and physiology. Lastly, our interpretation of the empirical results—in terms of evidence for top-down attention—needs to be applied with some caution, as we can only infer any attentional effects from the participants’ self-reports; and we can only assume that participants monitored their behaviour continuously. Future work could therefore use explicit measures of attention, perhaps supplemented by forms of external supervision and feedback, to validate behavioural effects.

Beyond this, our results open up a number of interesting questions for future research. It could be established whether the effects observed in our study can have long-lasting impact on the (generalizable) learning of motor control^68^. Another important question for future research is the potential attentional compensation of experimentally added sensory noise (e.g., via jittering or blurring the visual hand or via tendon vibration; although these manipulations may in themselves be ‘attention-grabbing’^69^). Finally, an interesting question is whether the observed effects could perhaps be reduced by actively ignoring or ‘dis-attending’^70,71^ away from vision. An analogous mechanism has been tentatively suggested by observed benefits of proprioceptive attenuation—thereby increasing the relative impact of visual information—during visuo-motor adaptation and visuo-proprioceptive recalibration^10,24,26,31,72-74^. These questions should best be addressed by combined behavioural and brain imaging experiments, to illuminate the neuronal correlates of the (supposedly attentional) precision weighting in the light of recently proposed implementations of predictive coding in the brain^37,40,75^.

To conclude, our results suggest a tight link between attention (precision control), multisensory integration, and action—allowing the brain to choose how much to rely on specific sensory cues to represent its body for action in a given context.

## Methods

### Task design

We used the same task design in the simulations and the behavioural experiment (see Fig. 1). For consistency, we will describe the task as performed by our human participants, but the same principles apply to the simulated agent. We designed our task as a non-spatial modification of a previously used hand-target tracking task^29,30^. The participant (or simulated agent) had to perform repetitive grasping movements paced by sinusoidal fluctuations in the size of a central fixation dot (sinusoidal oscillation at 0.5 Hz). Thus, this task was effectively a phase matching task, which we hoped to be less biased towards the visual modality due to a more abstract target quantity (oscillatory size change vs spatially moving target, as in previous studies). The fixation dot was chosen as the target to ensure that participants had to fixate the centre of the screen (and therefore look at the virtual hand) in all conditions. Participants (or the simulated agent) controlled a virtual hand model via a data glove worn on their unseen right hand (details below). In this way, vision (*seen* hand position via the virtual hand) could be decoupled from proprioception (*felt* hand position). In half of the movement trials, a temporal delay of 500 ms between visual and proprioceptive hand information was introduced by delaying vision (i.e., the seen hand movements) with respect to proprioception (i.e., the unseen hand movements performed by the participant or agent). In other words, the seen and felt hand positions were always incongruent (mismatching; i.e., phase-shifted) in these conditions. Crucially, the participant (agent) had to perform the hand-target phase matching task with one of two goals in mind: to match the target’s oscillatory phase with the seen virtual hand movements (vision) or with the unseen real hand movements (proprioception). This resulted in a 2 × 2 factorial design with the factors ‘visuo-proprioceptive congruence’ (congruent, incongruent) and ‘instructed modality’ (vision, proprioception).

### Predictive coding and active inference

We based our simulations on predictive coding formulations of active inference as situated within a free energy principle of brain function, which has been used in many previous publications to simulate perception and action^6,13,38,45,66,67^. Here, we briefly review the basic assumptions of this scheme (please see the above literature for details). Readers familiar with this topic should skip to the next section.

Hierarchical predictive coding rests on a probabilistic mapping of hidden causes to sensory consequences, as described by a hierarchical generative model, where each level of the model encodes conditional expectations (‘beliefs’; here, referring to subpersonal or non-propositional Bayesian beliefs in the sense of Bayesian belief updating and belief propagation; i.e., posterior probability densities) about states of the world that best explains states of affairs encoded at lower levels or—at the lowest level—sensory input. This hierarchy provides a deep model of how current sensory input is generated from causes in the environment; where increasingly higher-level beliefs represent increasingly abstract (i.e., hidden or latent) states of the environment. The generative model therefore maps from unobservable causes (hidden states) to observable consequences (sensory states). Model inversion corresponds to inferring the causes of sensations; i.e., mapping from consequences to causes. This inversion rests upon the minimisation of free energy or ‘surprise’ approximated in the form of prediction error. Model beliefs or expectations are thus updated to accommodate or ‘explain away’ ascending prediction error. This corresponds to Bayesian filtering or predictive coding^35-37^—which, under linear assumptions, is formally identical to linear quadratic control in motor control theory^76^.

Importantly, predictive coding can be implemented in a neurobiologically plausible fashion^6,35-37,40^. In such architectures, predictions may be encoded by the population activity of deep and superficial pyramidal cells, whereby descending connections convey predictions, suppressing activity in the hierarchical level below, and ascending connections return prediction error (i.e., sensory data not explained by descending predictions)^37,40^. Crucially, the ascending prediction errors are precision-weighted (where precision corresponds to the inverse variance), so that a prediction error that is afforded a greater precision has more impact on belief updating. This can be thought of as increasing the gain of superficial pyramidal cells^38,43^.

Active inference extends hierarchical predictive coding from the sensory to the motor domain; i.e., by equipping standard Bayesian filtering schemes (a.k.a. predictive coding) with classical reflex arcs that enable action (e.g., a hand movement) to fulfil predictions about hidden states of the world. In brief, desired movements are specified in terms of prior beliefs about state transitions (policies), which are then realised by action; i.e., by sampling or generating sensory data that provide evidence for those beliefs^38^. Thus, action is also driven by optimisation of the model via suppression of prediction error: movement occurs because high-level multi- or amodal prior beliefs about behaviour predict proprioceptive and exteroceptive (visual) states that would ensue if the movement was performed (e.g., a particular limb trajectory). Prediction error is then suppressed throughout a motor hierarchy; ranging from intentions and goals over kinematics to muscle activity^3,39^. At the lowest level of the hierarchy, spinal reflex arcs suppress proprioceptive prediction error by enacting the predicted movement, which also implicitly minimises exteroceptive prediction error; e.g. the predicted visual consequences of the action^5,33,40^. Thus, via embodied interaction with its environment, an agent can reduce its model’s free energy (‘surprise’ or, under specific assumptions, prediction error) and maximise Bayesian model evidence^77^.

Following the above notion of active inference, one can describe action and perception as the solution to coupled differential equations describing the dynamics of the real world (boldface) and the behaviour of an agent (italics)^6,38^.

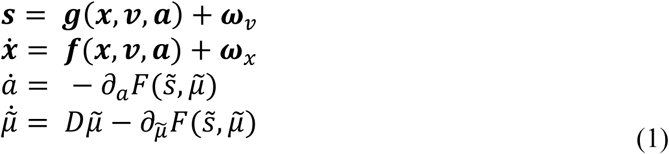

The first pair of coupled stochastic (i.e., subject to random fluctuations ***ω***_***x***_, ***ω***_***ν***_) differential equations describes the dynamics of hidden states and causes in the world and how they generate sensory states. Here, ***(s, x, ν, a)*** denote sensory input, hidden states, hidden causes and action in the real world, respectively. The second pair of equations corresponds to action and perception, respectively—they constitute a (generalised) gradient descent on variational free energy, known as an evidence bound in machine learning^78^. The differential equation describing perception corresponds to generalised filtering or predictive coding. The first term is a prediction based upon a differential operator *D* that returns the generalised motion of conditional (i.e., posterior) expectations about states of the world, including the motor plant (vector of velocity, acceleration, jerk, *etc*.). Here, the variables 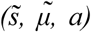correspond to generalised sensory input, conditional expectations and action, respectively. Generalised coordinates of motion, denoted by the ∼ notation, correspond to a vector representing the different orders of motion (position, velocity, acceleration, *etc*.) of a variable. The differential equations above are coupled because sensory states depend upon action through hidden states and causes ***(x, ν)*** while action *a(t)* = ***a(t)*** depends upon sensory states through internal states 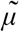. Neurobiologically, these equations can be considered to be implemented in terms of predictive coding; i.e., using prediction errors on the motion of hidden states—such as visual or proprioceptive cues about hand position—to update beliefs or expectations about the state of the lived world and embodied kinematics.

By explicitly separating hidden real-world states from the agent’s expectations as above, one can separate the generative process from the updating scheme that minimises free energy. To perform simulations using this scheme, one solves Eq. 1 to simulate (neuronal) dynamics that encode conditional expectations and ensuing action. The generative model thereby specifies a probability density function over sensory inputs and hidden states and causes, which is needed to define the free energy of sensory inputs:

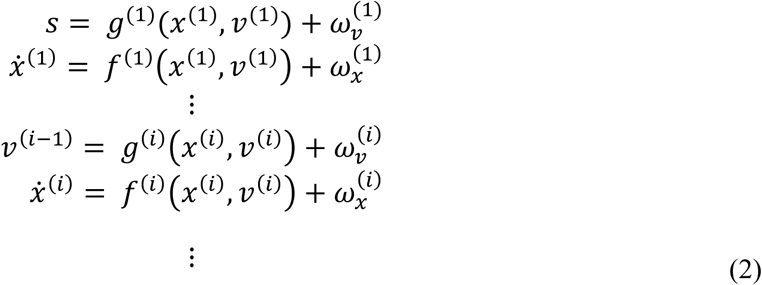

This probability density is specified in terms of nonlinear functions of hidden states and causes (*f*^*(i)*^, *g*^*(i)*^) that generate dynamics and sensory consequences, and Gaussian assumptions about random fluctuations (*ωx*^*(i)*^, *ω*_*ν*_^*(i)*^) on the motion of hidden states and causes. These play the role of sensory noise or uncertainty about states. The precisions of these fluctuations are quantified by (Π*x*^*(i)*^, Π_*ν*_^*(i)*^), which are the inverse of the respective covariance matrices.

Given the above form of the generative model (Eq. 2), we can now write down the differential equations (Eq. 1) describing neuronal dynamics in terms of prediction errors on the hidden causes and states as follows:

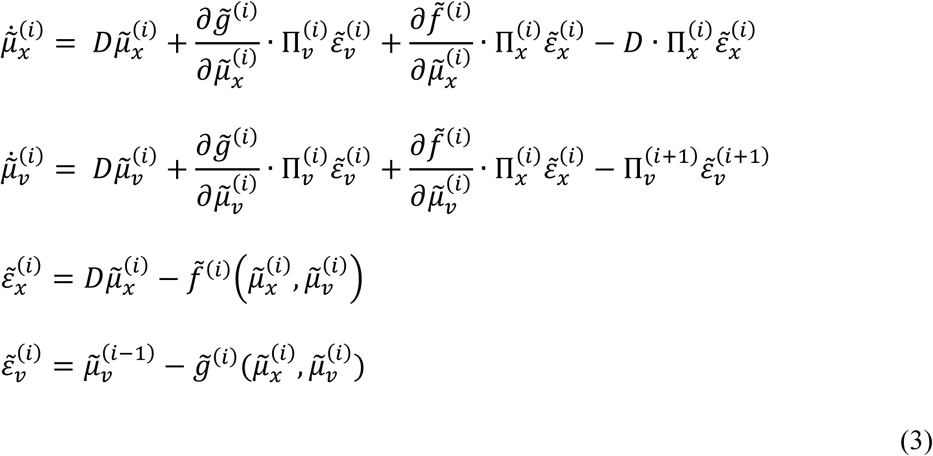

The above equation (Eq. 3) describes recurrent message passing between hierarchical levels to suppress free energy or prediction error (i.e., predictive coding^36,37^). Specifically, error units receive predictions from the same hierarchical level and the level above. Conversely, conditional expectations (‘beliefs’, encoded by the activity of state units) are driven by prediction errors from the same level and the level below. These constitute bottom-up and lateral messages that drive conditional expectations towards a better prediction to reduce the prediction error in the level below—this is the sort of belief updating described in the introduction.

Now we can add action as the specific sampling of predicted sensory inputs. As noted above, along active inference, high-level beliefs (conditional expectations) elicit action by sending predictions down the motor (proprioceptive) hierarchy to be unpacked into proprioceptive predictions at the level of (pontine) cranial nerve nuclei and spinal cord, which are then ‘quashed’ by enacting the predicted movements.

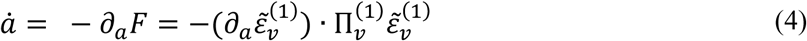

### Simulations of hand-target phase matching

In our case, the generative process and model used for simulating the target tracking task are straightforward (using just a single level) and can be expressed as follows:

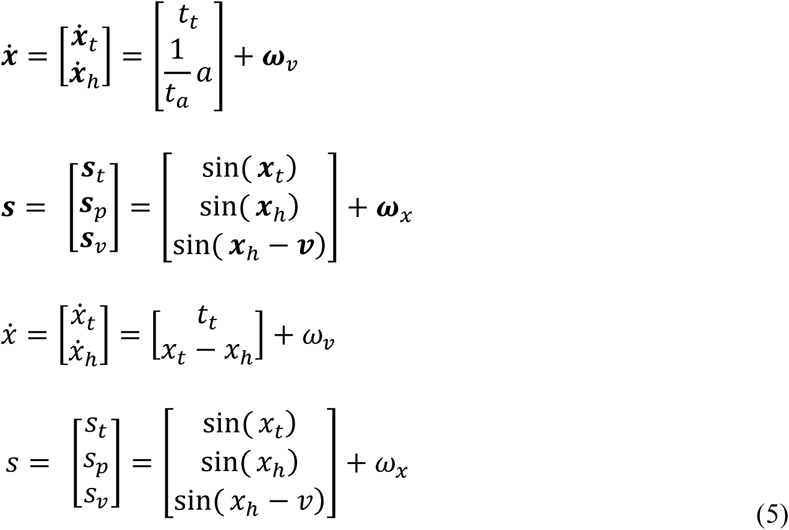

The first pair of equations describe the *generative process*; i.e., a noisy sensory mapping from hidden states and the equations of motion for states in the real world. In our case, the real-world variables comprised two hidden states ***x***_***t***_ (the state of the target) and ***x***_***h***_ (the state of the hand), which generate sensory inputs; i.e., proprioceptive ***s***_***p***_ and visual ***s***_***v***_ cues about hand posture, and visual cues about the target’s size ***s***_***t***_. Note that to simulate sinusoidal movements—as used in the experimental task—sensory cues pertaining to the target and hand are mapped via sine functions of the respective hidden states (plus random fluctuations). Both target and hand states change linearly over time, and become sinusoidal movements via the respective sensory mapping from causes to sensory data. We chose this solution in our particular case for a straightforward implementation of phase shifts (visuo-proprioceptive incongruence) via subtraction of a constant term from the respective sensory mapping (*v*, see below). Thus, the target state *x*_*t*_ is perturbed by hidden causes at a constant rate (*t*_*t*_ = 1/40), i.e., it linearly increases over time. This results in one oscillation of a sinusoidal trajectory via the sensory mapping sin(***x***_***t***_)—corresponding to one growing-and-shrinking of the fixation dot, as in the behavioural experiment—during 2 seconds (the simulations proceeded in time bins of 1/120 seconds, see Fig. 2). The hand state is driven by action *a* with a time constant of *t*_*a*_ = 16.67 ms, which induced a slight ‘sluggishness’ of movement mimicking delays in motor execution. Action thus describes the rate of change of hand posture along a linear trajectory—at a rate of 0.05 per time bin—which again becomes an oscillatory postural change (i.e., a grasping movement) via the sinusoidal sensory mapping. The hidden cause *v* modelled the displacement of proprioceptive and visual hand posture information in a virtue of being subtracted within the sinusoidal sensory mapping from the hidden hand state to visual sensory information sin(***x***_***t***_**-*v***). In other words, ***v*** = 0 when the virtual hand movements were congruent, and ***v*** = 0.35 (corresponding to about 111 ms delay) when the virtual hand’s movements were delayed with respect to the real hand. Note that random fluctuations in the process generating sensory input were suppressed by using high precisions on the errors of the sensory states and motion in the generative process (exp(16) = 8886110). This can be thought of as simulating the average response over multiple realizations of random inputs; i.e., the single movement we simulated in each condition stands in for the average over participant-specific realizations, in which the effects of random fluctuations are averaged out^38,66,67^. This ensured that our simulations reflect systematic differences depending on the parameter values chosen to reflect alterations of sensory attention via changing parameters of the agent’s model (as described below). The parameter values for the precision estimates are somewhat arbitrary and were adopted from previous studies using the same predictive coding formulation to simulate similar (oculomotor) tasks^38,66,67^. The key thing is that changing these parameters (i.e., the precision estimates for visual and proprioceptive cues) resulted in significant changes in simulated behaviour.

The second pair of equations describe the agent’s *generative model* of how sensations are generated using the form of Eq. 2. These define the free energy in Eq. 1 and specify behaviour (under active inference). The generative model has the same form as the generative process, with the important exceptions that there is no action and the state of the hand is driven by the displacement between the hand and the target *x*_*t*_ – *x*_*h*_. In other words, the agent believes that its grasping movements will follow the target’s oscillatory size change, which is itself driven by some unknown force at a constant rate (and thus producing an oscillatory trajectory as in the generative process). This effectively models (the compliance with) the task instruction, under the assumption that the agent already knows about the oscillatory phase of the target; i.e., it is ‘familiar with the task’. Importantly, this formulation models the ‘real hand’ instruction; under the ‘virtual hand’ instruction, the state of the hand was driven by *x*_*t*_ – (*x*_*h*_ – *v*), reflecting the fact that any perceived visual delay (i.e., the inferred displacement of vision from proprioception *v*) should now also be compensated to keep the virtual hand aligned with the target’s oscillatory phase under incongruence; the initial value for *v* was set to represent the respective information about visuo-proprioceptive congruence, i.e., 0 for congruent movement conditions and 0.35 for incongruent movement conditions. We defined the agent’s model to entertain a prior belief that visual and proprioceptive cues are normally congruent (or, for comparison, incongruent). This was implemented by setting the prior expectation of the cause *v* to 0 (indicating congruence of visual and proprioceptive hand posture information), with a log precision of 3 (corresponding to about 20.1). In other words, the hidden cause could vary, a priori, with a standard deviation of about exp(−3/2) = 0.22. This mimicked the strong association between seen and felt hand positions (under a minimal degree of flexibility), which is presumably formed over a lifetime and very hard to overcome and underwrites phenomena like the ‘rubber hand illusion’^16^ (see Introduction).

Crucially, the agent’s model included a precision-weighting of the sensory signals—as determined by the active deployment of attention along predictive coding accounts of active inference. This allowed us to manipulate the precision assigned to proprioceptive or visual prediction errors (Π_*p*_, Π_*v*_) that, per default, were given a log precision of 3 and 4, respectively (corresponding to 20.1 and 54.6, respectively). This reflects the fact that, in hand position estimation, vision is usually afforded a higher precision than proprioception^7,51^. To implement increases in task-related (selective) attention, we increased the log precision of prediction errors from the instructed modality (vision or proprioception) by 1 in each case (i.e., by a factor of about 2.7); in an alternative scenario, we tested for the effects of ‘incorrect’ allocation of attention to the non-instructed or ‘distractor’ modality by increasing the precision of the appropriate prediction errors. We did not simulate increases in both sensory precisions, because our study design was tailored to investigate selective attention as opposed to divided attention. Note that in the task employed, divided attention was precluded, since attentional set was induced via instructed task-relevance; i.e., attempted target phase-matching. In other words, under incongruence, only one modality could be matched to the target. The ensuing generative process and model are, of course, gross simplifications of a natural movement paradigm. However, this formulation is sufficient to solve the active inference scheme in Eq. 1 and examine the agent’s behaviour under the different task instructions and, more importantly, under varying degrees of selectively enhanced sensory precision afforded by an attentional set.

### Behavioural experiment

26 healthy, right-handed volunteers (15 female, mean age = 27 years, range = 19-37, all with normal or corrected-to-normal vision) participated in the experiment, after providing written informed consent. Two participants were unable to follow the task instructions during training and were excluded from the main experiment, resulting in a final sample size of 24. The experiment was approved by the local research ethics committee (University College London) and conducted in accordance with the usual guidelines.

During the experiment, participants sat at a table wearing an MR-compatible data glove (5DT Data Glove MRI, 1 sensor per finger, 8 bit flexure resolution per sensor, 60 Hz sampling rate) on their right hand, which was placed on their lap under the table. The data glove measured the participant’s finger flexion via sewn-in optical fibre cables; i.e., each sensor returned a value from 0 to 1 corresponding to minimum and maximum flexion of the respective finger. These raw data were fed to a photorealistic virtual right hand model^29,30^, whose fingers were thus moveable with one degree of freedom (i.e., flexion-extension) by the participant, in real-time. The virtual reality task environment was instantiated in the open-source 3D computer graphics software Blender (http://www.blender.org) using a Python programming interface, and presented on a computer screen at about 60 cm distance (1280 × 1024 pixels resolution).

The participants’ task was to perform repetitive right-hand grasping movements paced by the oscillatory size change of the central fixation dot, which continually decreased-and-increased in size sinusoidally (12 % size change) at a frequency of 0.5 Hz; i.e., this was effectively a phase matching task (Fig. 1). The participants had to follow the dot’s size changes with right-hand grasping movements; i.e., to close the hand when the dot shrunk and to open the hand when the dot grew. In half of the movement trials, an incongruence between visual and proprioceptive hand information was introduced by delaying the virtual hand’s movements by 500 ms with respect to the movements performed by the participant. The virtual hand and the real hand were persistently in mismatching postures in these conditions. The delay was clearly perceived by all participants.

Participants performed the task in trials of 32 seconds (16 movement cycles; the last movement was signalled by a brief disappearance of the fixation dot), separated by 6 second fixation-only periods. The task instructions (‘VIRTUAL’ / ‘REAL’) were presented before each respective movement trials for 2 seconds. Additionally, participants were informed whether in the upcoming trial the virtual hand’s movements would be synchronous (‘synch.’) or delayed (‘delay’). The instructions and the fixation dot in each task were coloured (pink or turquoise, counterbalanced across participants), to help participants remember the current task instruction, during each movement trial. Participants practised the task until they felt confident, and then completed two runs of 8 min length. Each of the four conditions ‘virtual hand task under congruence’ (VH cong), ‘virtual hand task under incongruence’ (VH incong), ‘real hand task under congruence’ (RH cong), and ‘real hand task under incongruence’ (RH incong) was presented 3 times per run, in randomized order.

To analyse the behavioural change in terms of deviation from the target (i.e., phase shift from the oscillatory size change), we averaged and normalized the movement trajectories in each condition for each participant (raw data were averaged over the four fingers, no further pre-processing was applied). We then calculated the phase shift as the average angular difference between the raw averaged movements of the virtual or real hand and the target’s oscillatory pulsation phase in each condition, using a continuous wavelet transform. The resulting phase shifts for each participant and condition were then entered into a 2 × 2 repeated measures ANOVA with the factors task (virtual hand, real hand) and congruence (congruent, incongruent) to test for statistically significant group-level differences. Post-hoc t-tests (two-tailed, with Bonferroni-corrected alpha levels to account for multiple comparisons) were used to compare experimental conditions.

After the experiment, participants were asked to indicate—for each of the four conditions separately— their answers to the following two questions: “How difficult did you find the task to perform in the following conditions?” (Q1, answered on a 7-point visual analogue scale from “very easy” to “very difficult”) and “On which hand did you focus your attention while performing the task?” (Q2, answered on a 7-point visual analogue scale from “I focused on my real hand” to “I focused on the virtual hand”). The questionnaire ratings were evaluated for statistically significant differences using a nonparametric Friedman’s test and Wilcoxon’s signed-rank test (with Bonferroni-corrected alpha levels to account for multiple comparisons) due to non-normal distribution of the residuals.

## Acknowledgements

We thank Thomas Parr for helpful comments on the simulations. This work was supported by funding from the European Union’s Horizon 2020 research and innovation programme under the Marie Skłodowska-Curie grant agreement No 749988 to JL. KF was funded by a Wellcome Trust Principal Research Fellowship (Ref: 088130/Z/09/Z).

## Supporting Information Legends

Raw data recorded during the two experimental runs for each participant. Each data file contains the real and virtual finger values (recorded by the data glove) alongside timing and experimental condition information. Please see the file ‘VariableCoding’ for details on variable names.

## References

1. Wolpert DM, Goodbody SJ, Husain M. Maintaining internal representations: the role of the human superior parietal lobe. Nature neuroscience. 1998;1(6):529.

2. Körding KP, Wolpert DM. Bayesian integration in sensorimotor learning. Nature. 2004;427(6971):244.

3. Kilner JM, Friston KJ, Frith CD. Predictive coding: an account of the mirror neuron system. Cognitive processing. 2007;8(3):159–166.

4. Shadmehr R, Krakauer JW. A computational neuroanatomy for motor control. Experimental brain research. 2008;185(3):359–381.

5. Friston K. What is optimal about motor control? Neuron. 2011;72:488–98.

6. Friston KJ, Daunizeau J, Kilner J, Kiebel SJ. Action and behavior: a free-energy formulation. Biological cybernetics. 2010;102(3):227–260.

7. Van Beers RJ, Sittig AC, Gon JJDVD. Integration of proprioceptive and visual position-information: An experimentally supported model. Journal of neurophysiology. 1999;81(3):1355–1364.

8. Van Beers RJ, Wolpert DM, Haggard P. When feeling is more important than seeing in sensorimotor adaptation. Current biology. 2002;12:834–837.

9. Foulkes AJM, Miall RC. Adaptation to visual feedback delays in a human manual tracking task. Experimental Brain Research. 2000;131(1):101–110.

10. Ingram HA, Van Donkelaar P, Cole J, Vercher JL, Gauthier GM, Miall RC. The role of proprioception and attention in a visuomotor adaptation task. Experimental Brain Research. 2000;132(1):114–126.

11. Sober SJ, Sabes PN. Flexible strategies for sensory integration during motor planning. Nature neuroscience. 2005;8(4):490.

12. Vijayakumar S, Hospedales T, Hait, A. Generative probabilistic modeling: understanding causal sensorimotor integration. Sensory Cue Integration. 2011;63–81.

13. Friston K. Prediction, perception and agency. International Journal of Psychophysiology. 2012;83(2):248–252.

14. Samad M, Chung AJ, Shams L. Perception of body ownership is driven by Bayesian sensory inference. PloS one. 2015;10:e0117178.

15. Rohe T, Noppeney U. Distinct computational principles govern multisensory integration in primary sensory and association cortices. Current Biology. 2016;26(4):509–514.

16. Botvinick M, Cohen J. Rubber hands ‘feel’touch that eyes see. Nature. 1998;391(6669):756.

17. Pavani F, Spence C, Driver J. Visual capture of touch: Out-of-the-body experiences with rubber gloves. Psychological science. 2000;11(5):353–359.

18. Holmes NP, Crozier G, Spence C. When mirrors lie:”Visual capture” of arm position impairs reaching performance. Cognitive, Affective, & Behavioral Neuroscience. 2004;4(2):193–200.

19. Holmes NP, Snijders HJ, Spence C. Reaching with alien limbs: Visual exposure to prosthetic hands in a mirror biases proprioception without accompanying illusions of ownership. Perception & Psychophysics. 2006;68(4):685–701.

20. Tsakiris M, Haggard P. The rubber hand illusion revisited: visuotactile integration and self-attribution. Journal of Experimental Psychology: Human Perception and Performance. 2005;31(1):80.

21. Makin TR, Holmes NP, Ehrsson HH. On the other hand: dummy hands and peripersonal space. Behavioural brain research. 2008;191(1):1–10.

22. Heed T, Gründler M, Rinkleib J, Rudzik FH, Collins T, Cooke E, O’Regan JK. Visual information and rubber hand embodiment differentially affect reach-to-grasp actions. Acta psychologica. 2011;138(1):263–271.

23. Limanowski J, Blankenburg F. Integration of visual and proprioceptive limb position information in human posterior parietal, premotor, and extrastriate cortex. Journal of Neuroscience. 2016;36(9):2582–2589.

24. Balslev D, Christensen LO, Lee JH, Law I, Paulson OB, Miall RC. Enhanced accuracy in novel mirror drawing after repetitive transcranial magnetic stimulation-induced proprioceptive deafferentation. Journal of Neuroscience. 2004;24(43):9698–9702.

25. Grafton ST, Schmitt P, Van Horn J, Diedrichsen J. Neural substrates of visuomotor learning based on improved feedback control and prediction. Neuroimage. 2008;39(3):1383–1395.

26. Bernier PM, Burle B, Vidal F, Hasbroucq T, Blouin J. Direct evidence for cortical suppression of somatosensory afferents during visuomotor adaptation. Cerebral Cortex. 2009;19(9):2106–2113.

27. Grefkes C, Ritzl A, Zilles K, Fink GR. Human medial intraparietal cortex subserves visuomotor coordinate transformation. Neuroimage. 2004;23:1494–1506.

28. Ogawa K, Inui T, Sugio T. Neural correlates of state estimation in visually guided movements: an event-related fMRI study. Cortex. 2007;43:289–300.

29. Limanowski J, Kirilina E, Blankenburg F. Neuronal correlates of continuous manual tracking under varying visual movement feedback in a virtual reality environment. Neuroimage. 2017;146:81–89.

30. Limanowski J, Friston K. Attentional modulation of vision vs proprioception during action. Cerebral Cortex. 2019:bhz192.

31. Taub E, Goldberg IA. Use of sensory recombination and somatosensory deafferentation techniques in the investigation of sensory-motor integration. Perception. 1974;3(4):393–408.

32. Friston K. The free-energy principle: a unified brain theory?. Nature reviews neuroscience. 2010;11(2):127.

33. Adams RA, Shipp S, Friston KJ. Predictions not commands: active inference in the motor system. Brain Structure and Function. 2013;218:611–643.

34. Vasser M, Vuillaume L, Cleeremans A, Aru J. Waving goodbye to contrast: self-generated hand movements attenuate visual sensitivity. Neuroscience of consciousness. 2019;1:niy013.

35. Rao RP, Ballard DH. Predictive coding in the visual cortex: a functional interpretation of some extra-classical receptive-field effects. Nature neuroscience. 1999;2(1):79.

36. Friston K, Kiebel S. Predictive coding under the free-energy principle. Philosophical Transactions of the Royal Society B: Biological Sciences. 2009;364(1521):1211–1221.

37. Bastos AM, Usrey WM, Adams RA, Mangun GR, Fries P, Friston KJ. Canonical microcircuits for predictive coding. Neuron. 2012;76(4):695–711.

38. Perrinet LU, Adams RA, Friston KJ. Active inference, eye movements and oculomotor delays. Biological cybernetics. 2014;108(6):777–801.

39. Grafton ST, Hamilton AFDC. Evidence for a distributed hierarchy of action representation in the brain. Human movement science. 2007;26(4):590–616.

40. Shipp S, Adams RA, Friston KJ. Reflections on agranular architecture: predictive coding in the motor cortex. Trends in Neuroscience. 2013;36:706–16.

41. Yon, D., & Press, C. (2017). Predicted action consequences are perceptually facilitated before cancellation. Journal of experimental psychology: human perception and performance, 43(6), 1073.

42. Limanowski J, Sarasso P, Blankenburg F. Different responses of the right superior temporal sulcus to visual movement feedback during self-generated vs. externally generated hand movements. European Journal of Neuroscience. 2018,47(4):314–320.

43. Feldman H, Friston K. Attention, uncertainty, and free-energy. Frontiers in human neuroscience. 2010;4:215.

44. Edwards MJ, Adams RA, Brown H, Parees I, Friston K. A Bayesian account of ‘hysteria’. Brain. 2012;135(11):3495–3512.

45. Brown H, Adams A, Parees I, Edwards M, Friston KJ. Active inference, sensory attenuation and illusions. Cognitive processing. 2013;14(4):411–427.

46. Brass M, Zysset S, von Cramon DY. The inhibition of imitative response tendencies. Neuroimage. 2011;14(6):1416–1423.

47. Kilner JM, Paulignan Y, Blakemore SJ. An interference effect of observed biological movement on action. Current biology. 2003;13(6):522–525.

48. Posner MI, Nissen MJ, Klein RM. Visual dominance: an information-processing account of its origins and significance. Psychological review. 1976;83(2):157.

49. Posner MI, Nissen MJ, Ogden WC. Attended and unattended processing modes: The role of set for spatial location. Modes of perceiving and processing information. 1978;137(158):2.

50. Rohe T, Noppeney U. Reliability-weighted integration of audiovisual signals can be modulated by top-down attention. Eneuro. 2018;5(1).

51. Kelso JA, Cook E, Olson ME, Epstein W. Allocation of attention and the locus of adaptation to displaced vision. Journal of Experimental Psychology: Human Perception and Performance. 1975;1(3):237.

52. Warren DH, Cleaves WT. Visual-proprioceptive interaction under large amounts of conflict. Journal of Experimental Psychology. 1971;90:206–214.

53. Redding GM, Clark SE, Wallace B. Attention and prism adaptation. Cognitive psychology. 1985;17(1):1–25.

54. Kelso JS. Motor-sensory feedback formulations: are we asking the right questions? Behavioral and Brain Sciences. 1979;2(1):72–73.

55. Foxe JJ, Simpson GV. Biasing the brain’s attentional set: II. Effects of selective intersensory attentional deployments on subsequent sensory processing. Experimental brain research. 2005;166(3-4):393–401.

56. Cressman EK, Henriques DY. Sensory recalibration of hand position following visuomotor adaptation. Journal of neurophysiology. 2009;102(6):3505–3518.

57. Rand MK, Heuer H. Visual and proprioceptive recalibrations after exposure to a visuomotor rotation. European Journal of Neuroscience. 2019.

58. Deneve S, Latham PE, Pouget A. Efficient computation and cue integration with noisy population codes. Nature neuroscience. 2001;4:826.

59. Ernst MO, Banks MS. Humans integrate visual and haptic information in a statistically optimal fashion. Nature. 2002;415:429.

60. Ernst MO. Learning to integrate arbitrary signals from vision and touch. Journal of Vision. 2007;7(5):7–7.

61. Ma WJ, Pouget A. Linking neurons to behavior in multisensory perception: A computational review. Brain research. 2008;1242:4–12.

62. Hospedales TM, Vijayakumar S. Structure inference for Bayesian multisensory scene understanding. IEEE transactions on pattern analysis and machine intelligence. 2008;30(12):2140–2157.

63. Kayser C, Shams L. Multisensory causal inference in the brain. PLoS biology. 2015;13:e1002075.

64. Körding KP, Beierholm U, Ma WJ, Quartz S, Tenenbaum JB, Shams L. Causal inference in multisensory perception. PLoS one. 2007;2:e943.

65. Vijayakumar S, Hospedales T, Hait, A. Generative probabilistic modeling: understanding causal sensorimotor integration. Sensory Cue Integration. 2011;63–81.

66. Adams RA, Aponte E, Marshall L, Friston KJ. Active inference and oculomotor pursuit: The dynamic causal modelling of eye movements. Journal of Neuroscience Methods. 2015;242:1–14.

67. Adams RA, Bauer M, Pinotsis D, Friston KJ. Dynamic causal modelling of eye movements during pursuit: confirming precision-encoding in V1 using MEG. Neuroimage. 2016;132:175–189.

68. Mathew J, Bernier PM, Danion FR. Asymmetrical relationship between prediction and control during visuo-motor adaptation. eNeuro. 2018;5(6).

69. Beauchamp MS, Pasalar S, Ro T. Neural substrates of reliability-weighted visual-tactile multisensory integration. Frontiers in Systems Neuroscience. 2010;4:25.

70. Clark A. Surfing uncertainty: Prediction, action, and the embodied mind. Oxford: Oxford University Press; 2015.

71. Limanowski J. (Dis-) attending to the Body: Action and Self-experience in the Active Inference Framework. In T. Metzinger & W. Wiese (Eds.), Philosophy and Predictive Processing. Frankfurt am Main: MIND Group; 2017.

72. Zeller D, Friston KJ, Classen JD. Dynamic causal modeling of touch-evoked potentials in the rubber hand illusion. Neuroimage. 2016;138:266–273.

73. Limanowski J, Blankenburg F. Network activity underlying the illusory self-attribution of a dummy arm. Human Brain Mapping. 2015a;36(6):2284–2304.

74. Limanowski J, Blankenburg F. That’s not quite me: limb ownership encoding in the brain. Social cognitive and affective neuroscience. 2015b;11(7):1130–1140.

75. Shipp S. Neural Elements for Predictive Coding. Frontiers in Psychology. 2016;7:1792.

76. Todorov E. General duality between optimal control and estimation. In 47th IEEE Conference on Decision and Control (pp. 4286–4292). IEEE; 2008.

77. Hohwy J. The Self-Evidencing Brain. Noûs. 2006;50:259–85.

78. Winn J, Bishop CM. Variational message passing. Journal of Machine Learning Research. 2005;6:661–94

